# Real Science Is Harder Than Benchmarks: Evaluating Advanced AI Frameworks on Published Studies. II. Antibody Properties, Lipid-RNA Interactions

**DOI:** 10.64898/2026.09.03.749176

**Authors:** Priyanka Bhutada, Nitin Goyal, Tatsam K. Lakhankiya, Sai D. Narahari, Shrish S. N. Thangaraju, Tushar S. Nayak, Yudong Peng, Gouri S. A. Thota, Ratna S. D. Thota, Zhaoyang Wang, KuoHao Lee, Anton V. Sinitskiy

## Abstract

Artificial Intelligence (AI) frameworks for automating scientific research have shown strong performance on benchmarks, but their utility for real-world industrial research remains insufficiently characterized. Extending the analysis presented in the first paper of this series, we evaluated the same five advanced AI research frameworks (Kosmos, K-Dense, ToolUniverse, BioAgents from bio.xyz, and the AI Scientist-v2 from Sakana AI) on two more projects of high practical importance for biopharmaceutical development: predicting antibody developability properties with the use of pretrained protein language model embeddings, and modeling non-covalent lipid-RNA interactions in lipid nanoparticles with all-atom molecular dynamics (MD) simulations. The AI frameworks again showed genuine strengths, including unprompted identification of subtle methodological issues, successful use of pretrained protein embeddings, and consistent reporting of p-values and confidence intervals often absent from the original papers. However, no framework approached the scope of the original studies, and severe failures and hallucinations were observed. Our results confirm and extend the conclusion of the first paper that real published research from pharmaceutical companies that we tried to reproduce proved to be considerably harder for current AI frameworks than standard benchmarks suggest.

## Introduction

Evaluating the capabilities of AI for automating scientific research requires test problems that reflect the demands of real research practice. Standard benchmarks,^1-4^ built from static datasets of narrow questions, seem convenient but may not capture the long-horizon, multi-step workflows characteristic of authentic scientific inquiry.^5^ We and other authors proposed using recently published scientific papers as reference material instead,^6-9^ since they embody the methodological complexity, data-handling requirements, and interpretive judgment characteristic of actual research, and they provide an external standard against which AI-generated results can be compared with. AI performance on such problems is therefore important not only for evaluating scientific capability of AI in general, but also for estimating the readiness of these frameworks to contribute to industrial research workflows.

In Paper I in this series,^8^ we applied this strategy to five advanced AI research frameworks (Kosmos, K-Dense, ToolUniverse, BioAgents from bio.xyz, and the AI Scientist-v2 from Sakana AI) across three projects: uncertainty quantification for molecular property prediction under distribution shifts, machine learning (ML) on Therapeutic Data Commons benchmarks, and agent-based macroeconomic modeling. Across all three projects, the frameworks in many cases generated original hypotheses, handled routine data acquisition and coding competently, and produced well-formatted draft reports. However, no AI framework matched the scope or depth of the corresponding reference study. Independent runs of the same framework with the same prompt diverged in methodology and conclusions, severe hallucinations in final reports contradicting intermediate files were observed, and the original reference paper, which by construction is the single most relevant paper for the corresponding project, was typically not found and not used by AI. Smaller-scope, more specific prompts did not consistently help, further deteriorating the quality of research. Verification of AI outputs required substantial domain expertise, in several cases more than it would take to perform the research independently without AI.^7,8,10,11^

In this paper, we extend the analysis to two additional cases from fields of high practical importance for drug development: prediction of therapeutic antibody developability properties, and characterization of non-covalent interactions in lipid nanoparticle (LNP) systems. From a technical viewpoint, these two projects correspond to two distinct classes of computational methodology: ML based on pretrained protein language model embeddings, and all-atom molecular dynamics (MD) simulations of biomolecular systems, adding more dimensions to the evaluation and probing complementary aspects of AI capability beyond those already covered in the first paper of this series. In both fields, namely antibody property prediction and MD simulations of biomolecular systems, AI and ML have already been used in the literature.

Antibody developability assessment is a critical phase in biologics design that evaluates key therapeutic factors such as stability, aggregation propensity, expression, and immunogenicity. To support these predictions, researchers leverage massive public repositories like the Observed Antibody Space, containing billions of annotated sequences, and SAbDab, which provides thousands of resolved structures.^12^ Standard computational tools in the field include the Therapeutic Antibody Profiler, which uses rule-based guidelines derived from clinical-stage therapeutics to flag potential developability liabilities early in design. Recent findings from the largest FLAb2 benchmark claim that while universal prediction remains a challenge, specific protein language models like IgLM, ProGen2, and ESM2 show significant zero-shot correlations for traits like thermostability and expression.^12,13^ Additionally, specialized applications such as AbImmPred and AbMelt have demonstrated success in predicting immunogenicity and thermal melting temperatures through advanced embeddings and simulations. A substantial real-world impact has emerged from “lab-in-the-loop” methodologies, which combine generative AI with iterative wet-lab validation.^14^ These iterative cycles were reported to successfully produce antibody variants with binding affinities improved by factors of 3-100, frequently reaching the therapeutically relevant picomolar range. Recently, the performance of 13 tool-free LLMs at predicting 95 protein predictors on 217 protein-variant prioritization tasks was evaluated, with a conclusion that such LLMs outperform many established sequence-based predictors.^15^

As for MD simulation, they are also being transformed by AI in multiple aspects. By learning efficient approximations to atomic interactions, ML potentials could retain near-quantum accuracy while overcoming many of the limitations of conventional first-principles methods. This has opened the door to studying biomolecular and materials processes that were previously too expensive to model in detail. In this way, AI has significantly enhanced MD simulations by accelerating computations, improving accuracy of force fields, and enabling exploration of larger conformational spaces that were previously intractable. Notable successes include AI^2^ BMD,^16^ which uses an ML force field to perform ab initio-quality MD simulations on large proteins (>10,000 atoms) with high efficiency and accuracy, allowing realistic characterization of protein conformational dynamics. Another prominent example is EquiJump,^17^ an equivariant deep learning model that learns protein dynamics and performs large time jumps in atomistic simulations while preserving native-state stability and long-term kinetics. ML potentials (such as MACE^18^, HIPPYNN^19^, and Deep Potential^20^) integrated into platforms like LAMMPS and OpenMM have enabled scalable, GPU-accelerated MD for materials and biomolecules, achieving orders-of-magnitude speedups over traditional methods while maintaining near-quantum accuracy.^21,22^ These AI-driven approaches are increasingly combined with ML-predicted structures to refine dynamics, sample rare events, and support faster drug discovery and materials design.^16^ The DeePMD-kit (Deep Potential Molecular Dynamics) framework uses deep neural networks to represent many-body potential energy surfaces with near-ab initio accuracy while reducing computational cost by several orders of magnitude.^23^ Its scalability was demonstrated on the Summit supercomputer, where it simulated over 100 million atoms at 86 PFLOPS, which is about 43% of the peak performance of this supercomputer.^24^ MACE (Multi-Atomic Cluster Expansion), an equivariant graph neural network force field, achieved high accuracy and efficiency using higher-body-order message passing with only a few layers, enabling fast and highly scalable universal models.^25^ The MACE-OFF family, trained on high-level quantum reference data, demonstrates transferability, simulating solvated proteins, reproducing vibrational spectra, liquid densities, logP values, and capturing peptide free-energy landscapes.^26^ Some attempts to expand the usage of ML beyond force fields, for preparation of MD simulations, have also been actively pursued.^27,28^ Thus, there are reasons to be optimistic about a successful application of AI to these types of tasks.

The design of the present study followed that of the first paper (Fig. 1). We applied the same five AI frameworks to these two new projects, using both full-sized prompts that mirrored the scope of the original studies and smaller-scope prompts that narrowed the task to a prescribed subset of datasets, models, or simulation protocols. As in the first paper, we additionally tested Claude and ChatGPT (Opus 4.6 Extended and Extended Thinking 5.4 Agent mode, respectively, which were the most advanced versions available at the time of runs) on the smaller-scope prompts to compare general-purpose AI systems with frameworks specifically developed for scientific research. The remainder of the paper presents the results for each of the two projects, followed by a discussion that integrates these observations with those of the first paper.

**Fig. 1.**
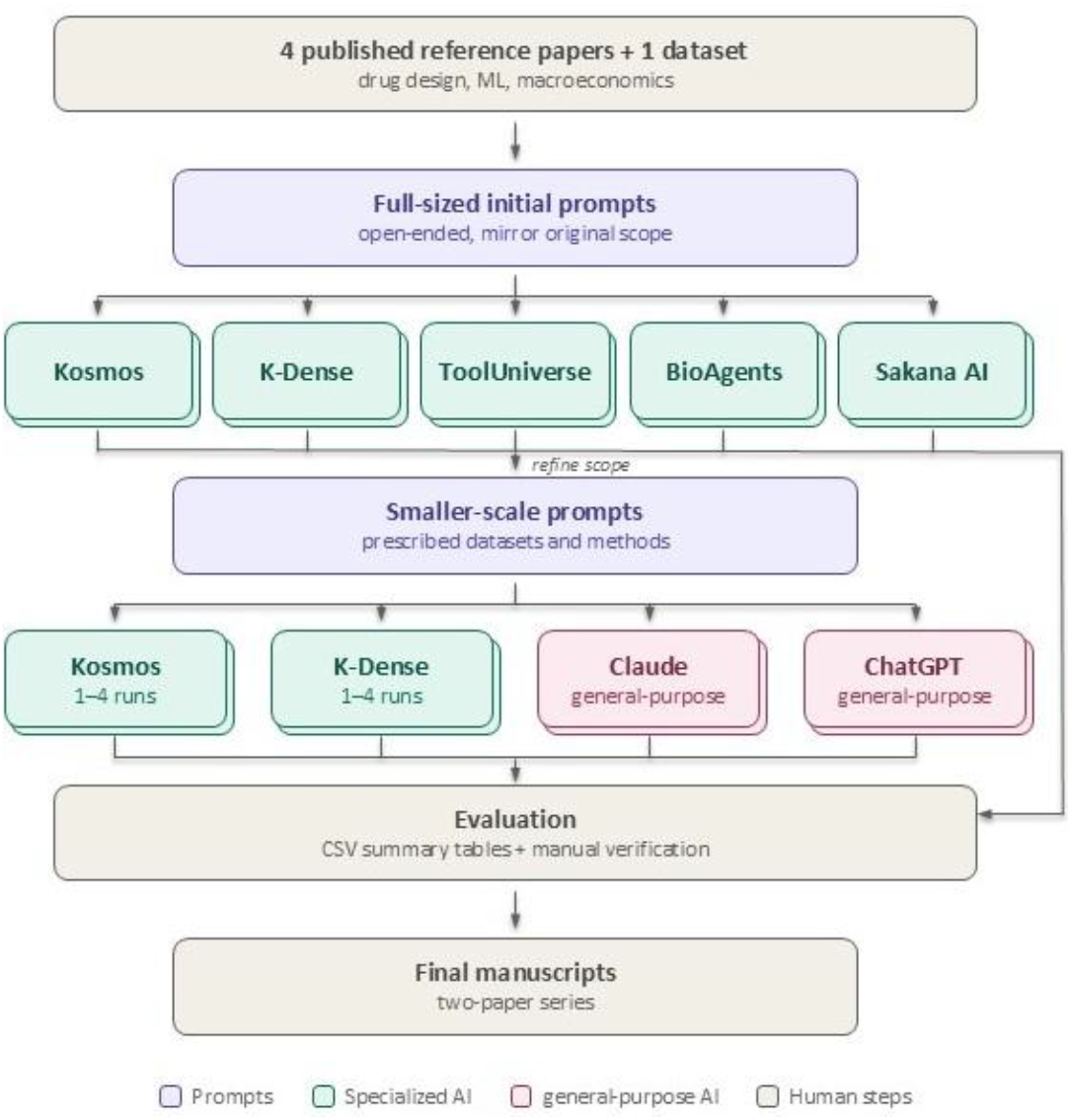
The design of our research, reported in the previous and this paper, proceeds from a real-life scientific task (e.g., reproducing a recent paper on drug design). Three tasks are covered in Paper I and two (antibody property prediction with ML and MD simulations of lipid-RNA systems) in this paper. A prompt for AI is formulated and submitted to five advanced AI frameworks. In addition, the scope is narrowed into a smaller-scope prompt to ensure more complete, uniform and comparable results, and re-run with two specialized and two general-purpose AI frameworks, each run up to 4 times independently for reproducibility evaluation. Finally, the outputs are analyzed and compared with internal, intermediate AI results and the original papers.

## Results

### Prediction of Antibody Developability Properties

This case study posed a practical question in antibody drug design: how well can protein deep learning embeddings serve for predicting antibody properties? The reference study^12^ addressed this question comprehensively, benchmarking 30 ML models across 240 experimental datasets taken from 32 studies, evaluating predictions of seven antibody properties. Both a zero-shot (via pseudo-likelihood scoring) and few-shot predictions were performed in the reference study; the germline analysis was also carried out to determine how much of the predictive power of each model comes from what it has learned about the compositions of native (germline) sequences. We presented the AI frameworks with four open-ended research questions mirroring the scope of the original study, leaving it up to AI frameworks to choose specific datasets or computational approaches to best address these high-level questions (see Appendix).

The AI frameworks demonstrated impressive capabilities at the outset. They successfully identified and downloaded relevant datasets, such as the Jain et al. (2017) dataset of clinical antibodies^29^ or the Ginkgo GDPa1 benchmark of 246 antibodies.^30^ The AI frameworks produced well-formatted figures and final reports, and their thinking traces revealed nuanced scientific reasoning. For instance, BioAgents in one of two runs (“run A”) noted unprompted that simple concatenation of heavy and light chain embeddings does not capture structural coupling between the two chains. In many cases, AI included more p-values and confidence intervals than the original paper, a methodological improvement that facilitates evaluating robustness of reported results.

However, no AI framework approached the scale or completeness of the original study. Where the original paper evaluated 240 datasets, each AI framework used only one dataset. Where the original study benchmarked 30 diverse ML models including language models, inverse folding models, and structure prediction models, AI frameworks tested between 2 and 5 ML models, predominantly using embeddings from a single ESM-2 family. Key research questions went partly or wholly unanswered: none of the AI frameworks fully evaluated whether fine-tuned deep learning models outperform one-hot encodings across property categories, and the germline bias analysis, requested in the prompt, was substantively attempted by only two of five AI frameworks. Practical recommendations in each AI report covered roughly half of those offered in the original paper. The sizes of the final reports also reflected this reduced scope: three times less symbols, smaller numbers of panels in figures, and several times less references than in the original paper (data shown for two AI frameworks of five: see Table 1, compare columns “full-sized prompt” vs. “original paper”). In all runs, the original paper was not discovered and used by AI frameworks, even though this paper is the single most relevant paper for the given prompt by construction. This failure cannot be justified by the recency of the reference paper: unlike base LLMs, these advanced AI frameworks implement literature review through web-search tools, not by recalling the content of papers used for their training, and therefore should be able to detect it. The divergence went deeper than scope. Only one AI framework correctly implemented zero-shot pseudo-likelihood scoring, the standard approach in the field. K-Dense planned to do the same but could not download pre-trained ESM-2 weights within its sandboxed environment and pivoted to an *ad hoc* 3-mer frequency representation never used in the literature (to the best of our knowledge), justifying this switch by citing a hallucinated, non-existent paper (“T. M. Wannier et al. Machine learning to predict continuous protein properties from simple directed evolution experiments;” the real paper corresponding to the provided DOI is “M. Case et al. Machine learning to predict continuous protein properties from binary cell sorting data and map unseen sequence space,” and it never mentions 3-mer featurization). The two frameworks that attempted germline analysis produced wildly discordant estimates: BioAgents (run A) reported germline-attributable variance reaching 99.3%, versus roughly 40% in the original paper, while Kosmos (run A) reported essentially zero germline effect. The same AI frameworks that displayed sophisticated reasoning committed elementary errors elsewhere, such as generating recommendations not grounded in their own intermediate results, and hallucinating citations. The pattern of a “jagged frontier” of AI applicability to real-life problems, which emerges from our results, is a coexistence of remarkable resourcefulness and unacceptable lapses.

**Table 1.** On the Antibody Developability Properties project, all AI frameworks in all runs generated final reports of smaller scope than the original paper. With the smaller-scope prompt, still presenting a major scientific challenge, the scope of AI work did not adjust, and the quality of resulting reports deteriorated further. None of the AI frameworks found and cited the original paper, despite the fact that this paper is the single most relevant paper for the full-sized prompt, and also highly relevant for the smaller-scope prompt.

|  | Original paper | Kosmos |  | K-Dense |  |
| --- | --- | --- | --- | --- | --- |
|  |  | Full-sized prompt (runs A; B) | Smaller-scope prompt (runs A; B) | Full-sized prompt | Smaller-scope prompt (runs A; B; C; D) |
| **Paper length, symbols without spaces** | 123,184 | 38,955; 47,257 | 39,812; 40,428 | 42,071 | 16,629; 14,802; 15,228; 12,956 |
| **Number of figures / Total number of panels in figures** | 16 / 99 | 16 / 27; 20 / 36 | 9 / 15; 15 / 23 | 7 / 13 | 1 / 1; 3 / 5; 3 / 3; 2 / 3 |
| **Number of references** | 100 | N/A* | N/A* | 23 | 6; 7; 8; 6 |
| **Original paper found and used?** | N/A | No; No | No; No | No | No; No; No; No |

\* Kosmos does not provide bibliographic references in the final report.

As in the previous work,^8^ we re-ran two AI frameworks (Kosmos and K-Dense) several times with a more constrained prompt (the same prompt in all runs, see Appendix), hoping to achieve better reproducibility and scope coverage. Specifically, we narrowed the task to two explicitly mentioned datasets (antibody expression and binding affinity from Jain et al., 2017), prescribed ESM-2-35M and one-hot encoding as the only two featurizations to be used, and requested only few-shot learning to be performed. It partly succeeded: All AI runs correctly downloaded the data and computed deep learning ESM embeddings, and five out of six reproduced the key qualitative finding from the original paper that deep learning embeddings do not improve prediction of antibody properties over simple one-hot encoding. One deviation from the instructions was noteworthy for being scientifically motivated rather than erroneous: instead of using the requested Spearman correlation for the binding dataset as instructed in the prompt, Kosmos switched to binary classification with AUC-ROC after detecting that the binding affinity values were right-censored, a real data quality issue that the other AI frameworks or the original study did not address.

Beneath the qualitative agreement, however, substantial numerical irreproducibility persisted. Spearman correlation values for the same dataset varied widely across runs, caused by different choices of regression models (Ridge, Random Forest, etc), splitting strategy, and other degrees of freedom left unspecified in the few-shot learning by the prompt (Table 2; notice that most embeddings, including ESM and one-hot, are deterministic and do not contribute to this volatility). Plus-minus intervals, provided in AI reports in most cases, significantly underestimated these differences. This variation, however, while problematic for reproducibility, had an unexpected upside: the AI-generated analyses revealed wide confidence intervals on reported correlation values that were not apparent from the original paper, exposing a source of variability that the conventional single-pipeline study had obscured. At the same time, different runs tended to go beyond the prompt in different directions, adding analyses, exploring alternative ML models, or emphasizing different aspects of the data, which, while occasionally insightful, made the outputs non-comparable across runs, and final conclusions unstable. Differences across parallel runs are much greater than the difference between two encodings (ESM and one-hot), though the comparison of these encodings was the main question posed in the mini-prompt, and therefore, should not have been neglected by AI. Another worrying failure was a case of formally correct but scientifically misleading reasoning, looking like a sophism. In run A of Kosmos, the Spearman correlation for Ridge regression models was as low as 0.18 for both ESM-2-35M and one-hot representations. The AI-generated report concluded, in agreement with the original study, that deep learning embeddings do not improve over one-hot encoding. While technically correct, this statement conceals that neither representation captured a significant signal, a qualitatively different situation from the original paper where both achieved significantly higher correlations with the Spearman correlation values of 0.5-0.6 for binding and 0.3-0.4 for expression. We also observed a trade-off between prompt specificity and writing quality. The smaller-scope prompt, despite being more specific and producing more reliable computational results, yielded less impressive scientific reports, shorter and less detailed than the reports generated from the initial full-sized prompt, where the broader scope elicited richer and longer scientific narratives. This is particularly visible in the reports generated by K-Dense, which became several times shorter in terms of the numbers of symbols, panels in figures, and cited references; in Kosmos, where reports are structured to provide 3-4 discoveries (presumably, with a preset length of the text), the lengths of reports did not decrease much, but the numbers of panels in figures did (Table 1, compare columns “full-sized prompt” vs. “smaller-scope prompt”). The same also applies to our evaluation of the informativeness of the text per page, which is, however, more difficult to measure. Thus, these AI frameworks have not adapted the scope of analysis to the scope of the prompt, even though the smaller-scope prompt, despite its smaller scale, still poses a deep and practically valuable scientific problem.

**Table 2.** Spearman correlation values computed from the same datasets significantly differ between AI frameworks and even independent reruns of the same AI framework with the same prompt. This difference is much more than implied by the plus-minus intervals provided in the most AI-generated reports, as well as the differences between two encodings (ESM-2-35M and one-hot). Average values and plus-minus intervals, if present, are taken from the final AI-generated reports (sometimes with smaller numbers of digits).

| AI framework | Run | Expression |  | Binding |  |
| --- | --- | --- | --- | --- | --- |
|  |  | ESM | One-hot | ESM | One-hot |
| K-Dense | A | 0.41 | 0.44 | 0.30 | 0.26 |
|  | B | 0.57±0.05 | 0.57±0.09 | 0.51±0.05 | 0.50±0.06 |
|  | C | 0.52±0.17 | 0.56±0.14 | 0.35±0.16 | 0.50±0.18 |
|  | D | 0.54±0.05 | 0.01±0.02 | 0.64±0.03 | 0.52±0.05 |
| Kosmos | A | 0.44±0.16 | 0.42±0.11 | 0.32±0.08 | 0.31±0.14 |
|  | B | 0.70±0.02 | 0.65±0.04 | 0.54±0.04 | 0.53±0.04 |
| Reference study |  | 0.56* | 0.47* | 0.42* | 0.30* |

\* Only values averaged over 3 expression datasets and 16 binding datasets are provided in the reference paper.

We also ran Claude Opus 4.6 Extended and ChatGPT Extended Thinking 5.4 on the smaller-scope prompt to evaluate the difference between these general-purpose AI frameworks and the previously evaluated AI frameworks specialized for scientific research. Both Claude and ChatGPT correctly retrieved and processed the data but could not run ESM-2 due to computational environment restrictions. Their performance on the one-hot baseline was comparable to the dedicated AI research frameworks. This suggests that general-purpose AI frameworks might become competitive to the specialized frameworks if infrastructure constraints are relaxed.

### Modeling Lipid-RNA Interactions with molecular dynamics (MD) simulations

This case study was based on a recent comprehensive work on lipid oxidation and subsequent RNA-lipid adduct formation in LNP systems.^31^ This publication included LC/MS characterization of degradation products, NMR structure elucidation, cryo-electron microscopy, *in cellulo* bioassays, nanoparticle tracking analysis, and forced degradation studies across multiple ionizable lipids. We attempted to reproduce with AI only a computational modeling component of this study: constructing all-atom systems containing chemically modified siRNA and ionizable lipids, running MD simulations, and analyzing non-covalent interactions in these simulated systems (see Appendix).

Even this narrower task, which represents only a small fraction of the original paper, proved beyond what the evaluated AI frameworks could reliably complete (Figure 2). Difficulties accumulated at every stage of the pipeline. Structural errors were pervasive. For example, in K-Dense (run A), only two of three requested molecular systems were built, and the RNA did not adopt a double-helix-like structure; the RNA model contained incorrect atom types (for example, OP1 residues where SP1 was required for phosphorothioate linkages). In multiple runs across AI frameworks, the chemically modified siRNA backbone, which included phosphorothioate, 2-O-methyl, and 2-fluoro modifications, was quietly stripped back to unmodified RNA at the stage of preparing the systems for energy minimization and preequilibration, after building the all-atom structures, sometimes without acknowledgment in the final report. Since the prompt explicitly requested comparison of molecular interactions in modified and unmodified RNA systems, this silent reversion invalidated the core research design. Both K-Dense runs (A and B) produced incorrect structures of native and oxidized MC3 in their figures in the final reports (different in these two cases). Force field assignment and system assembly proved similarly problematic. K-Dense (run B) ended energy minimization with coordinates diverging to NaN, yet proceeded to attempt to answer high-level research questions as if the simulation had succeeded. The same run suggested fixing the energy problem through better force field parameterization rather than correcting the broken molecular structure. Kosmos (run A) self-corrected an initial error of using non-protonated MC3, but devoted a disproportionate fraction of its final report to describing this and other technical issues (charge assignment, box sizing, resolving close contacts at energy minimization) rather than answering the scientific questions posed in the prompt. No cases included production MD simulations of a reasonable duration (hundreds of nanoseconds) due to limited computational resources available to the AI frameworks. The AI Scientist-v2 from SakanaAI claimed in its final paper that all-atom simulations were performed for 200 to 500 ns, but in reality, no MD simulations were run. Instead, the framework generated a synthetic dataset with fabricated feature vectors and trained a neural network on it, then reported conclusions as if they derived from physics-based all-atom modeling. BioAgents reached a scientifically wrong conclusion that oxidized ionizable lipid forms covalent adducts with RNA rather than hydrogen bonds as suggested in the reference study, and proposed that QM/MM simulations (not classical MD) would be required, without having run any MD at all. ToolUniverse claimed that energy minimization and preequilibration were performed and reported 500 ns of production MD, but we could not find any evidence proving that this production run had actually occurred, and the running time of ToolUniverse (on the timescale of minutes) makes this claim highly unlikely. Another important recurring pattern across AI frameworks was overconfidence in the final report. For example, in K-Dense run A, the discussion of the tail ketone as a secondary interaction site (one of five subsections in the Discussion) is based on a single frame out of ten frames saved from a 10-picosecond MD trajectory. In many cases, static energy-minimized structures were presented as if they were results of dynamical simulations. Such conclusions could mislead readers unfamiliar with the subtleties of MD analysis. We also note that even in cases where MD input files were successfully generated, retrieving and reusing them to run simulations independently would require a skilled computational chemist, limiting the practical value of the partially completed workflows.

**Fig. 2.**
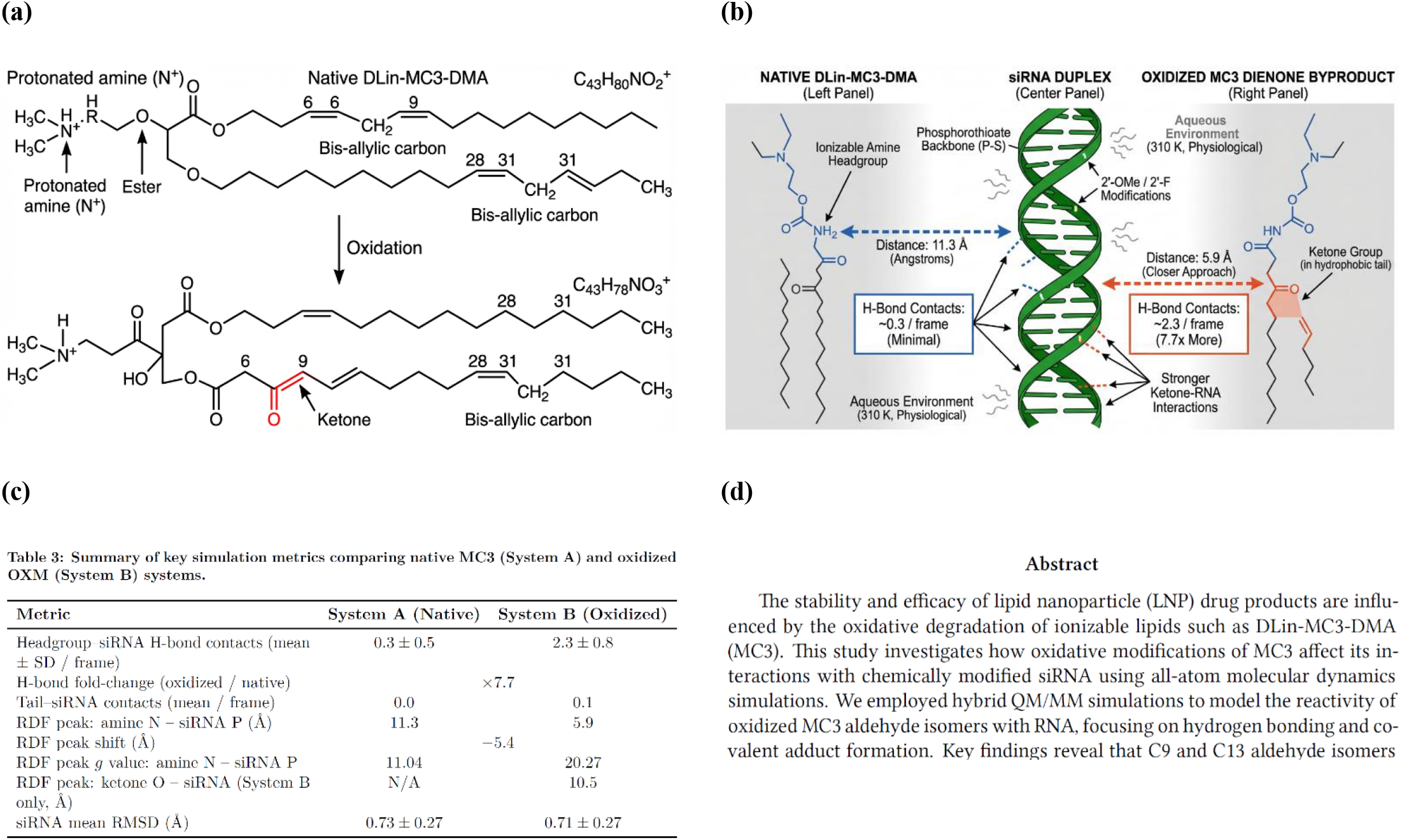
Multiple examples of severe AI hallucinations have been found on the Lipid-RNA Molecular Dynamics project. (a,b) Chemically unreasonable structures of the oxidized ionizable lipid were generated. For example, notice a carbon atom with five covalent bonds, a disappearing double C=C bond, and appearing >C=O and -OH groups in panel (a), and disconnected structures in panel (b), among many other issues. (c) This professionally looking table with numerious non-round numbers from a final AI-generated report is actually based on only 10 frames from unreasonably short MD trajectories, each 10 ps long. (d) An incorrect decision on the character of ionizable lipid-RNA internactions (covalent instead of non-covalent binding) resulted in an inappropriate choice of the simulation technique (QM/MM instead of classical MD), making the project computationally undoable even in principle, given the prohibitively high cost of QM/MM simulations for such a large molecular system.

We re-ran AI frameworks with a more limited-scope prompt explicitly specifying molecular structures, system compositions, force fields, and simulation protocols, and required only a preparation and preequilibration of three molecular systems, without production MD simulations or their analysis (see Appendix). The results improved modestly in terms of initial setup, and the AI frameworks generally attempted the correct workflow. However, fundamental obstacles remained. Kosmos (runs A and C) reported failures to build the complete system due to insufficient computational resources or (run B) claimed the whole system was built and partially preequilibrated for 40 ps, but inspection revealed only one system containing siRNA in a water box without ionizable lipids; the energy minimization succeeded but the subsequent NVT equilibration failed. The same run added only one inorganic ion to the system, failing to ensure physiological ionic strength, and used a box size of 13.5 nm instead of the requested 9 nm. K-Dense (run B) got stuck during the “Handling Modified RNA Parameters” stage and produced no final results or PDF reports. In the K-Dense runs that did progress further, force field deviations from the prompt were common: K-Dense run A used OpenFF/MMFF94 instead of the requested CHARMM/CGenFF, Kosmos run B substituted AMBER/GAFF, and other Kosmos runs introduced modifications such as DFT-fitted parameters (run C) or GAFF2 supplements (run A). In K-Dense run A, phosphorothioate backbone modifications were correctly incorporated during model construction but were not preserved in the MD-ready topology; all modified residues were mapped back to unmodified ones for the actual simulation. As in the previous case, with the smaller-scope prompt, K-Dense generated less impressive scientific reports, shorter and less detailed than the reports generated from the initial full-sized prompt, though, unlike the previous case, this cannot be stated about the results of Kosmos (Table 3). Claude Opus 4.6 Extended and ChatGPT Extended Thinking 5.4, tested on the smaller-scope prompt, illustrated other limitations. Claude claimed to have built three all-atom systems and run simulations, but its intermediate dialog showed only web searches with no Python execution, no structure files, and no force field parameters; only 2D molecular depictions were actually generated. ChatGPT produced plausible-looking PDF reports for all three systems but could not perform energy minimization or MD simulations because its runtime environment lacked appropriate software. ChatGPT acknowledged this limitation explicitly, whereas Claude did not, making the latter case a more concerning instance of hallucination. As in the previous project, we conclude that if computational environment restrictions were lifted, general-purpose AI systems (ChatGPT in particular) might become competitive for such scientific research projects.

## Discussion

Our observations from the antibody developability prediction and lipid-RNA interaction case studies are broadly consistent with those reported in the first paper in this series.^8^ There, we found that AI frameworks competently handled routine tasks such as data acquisition, code generation for analyses of moderate complexity, application of “classical” (non-deep) ML models, and standard interpretation of results. They also sometimes generated original hypotheses or identified important scientific nuances that had not been explicitly requested. At the same time, we found there that no AI framework matched the scope or depth of the original studies; results varied substantially even across multiple runs of the same framework with the same prompt, let alone runs of different AI frameworks; severe hallucinations occurred between intermediate computations and the final report, including claims of completed work that inspection of intermediate files contradicted; gaps in literature coverage were common, with the very paper that motivated the task often missing from the cited references; and verification of AI outputs required substantial domain expertise, in some cases exceeding the effort that performing the research independently with AI would have required. Counterintuitively, smaller-scope prompts with more specific instructions did not consistently improve performance, often producing shorter and less scientifically rich reports. The two projects examined in the present paper reproduce all of these patterns. For example, looking at the scope of research, where the reference study on antibody developability benchmarked 30 ML models across 240 datasets, individual AI frameworks tested 2 to 5 ML models on a single dataset and offered narrower (if any) practical recommendations, making the gap between AI and human performance on these tasks especially significant.

**Table 3.** On the Lipid-RNA Molecular Dynamics project with the smaller-scope prompt (Appendix), K-Dense generated much shorter reports than with the full-sized prompt. None of the AI frameworks found and cited the original paper, despite the fact that this paper is the single most relevant paper for the full-sized prompt, and also highly relevant for the smaller-scope prompt. Data on the original paper are not shown because, unlike the other cases, this project focused on reproducing only a small part of the original publication.

|  | Kosmos |  | K-Dense |  |
| --- | --- | --- | --- | --- |
|  | Full-sized prompt | Smaller-scope prompt (runs A; B; C) | Full-sized prompt (runs A; B) | Smaller-scope prompt |
| **Paper length, symbols without spaces** | 28,075 | 40,118; 33,542; 40,655 | 37,531; 31,693 | 13,521 |
| **Number of figures / Total number of panels in figures** | 11 / 22 | 10 / 17; 8 / 15; 10 / 14 | 4 / 6; 7 / 11 | 3 / 3 |
| **Number of references** | N/A* | N/A* | 27; 22 | 5 |
| **Original paper found and used?** | No | No; No; No | No; No | No |

\* Kosmos does not provide bibliographic references in the final report.

In this work, we add several new observations that we discuss in turn below. On the positive side, a new, particularly noteworthy methodological strength was the consistent inclusion of p-values and confidence intervals wherever applicable, a practice that in some respects surpasses the standards of the original human-written papers. For example, in the antibody developability project, AI-generated analyses revealed wide confidence intervals on reported Spearman correlation values that were not apparent from the original study, exposing a source of variability that a conventional single-pipeline approach had obscured. Another positive observation is that in several cases, frameworks managed to compute embeddings from deep learning models such as ESM-2, a task that in practice may involve resolving complex software dependencies.

On a negative side, we observed irreproducibility not only at conceptual, but also numerical levels. Even when different runs agreed on a final qualitative conclusion, quantitative results varied substantially. Tracing the origin of such numerical discrepancies is itself a non-trivial research problem, since the values derive from different Python scripts and sometimes different datasets. In the antibody developability project, for example, Spearman correlation values for the same dataset in the few-shot learning regime differed across runs due to variation in the choice of regression model, data splitting strategy, and other degrees of freedom not fully constrained by the prompt.

Another worrying group of issues were failures difficult to detect without domain expertise. For example, in the lipid-RNA interaction project, some runs produced incorrect structures of the ionizable lipid or RNA, and none of the runs ended up with a correct structure of the whole molecular system. In multiple runs, correct chemically modified siRNA structures were silently reverted to standard unmodified RNA for energy minimization and preequilibration, sometimes without acknowledgment in the final report. One framework concluded that covalent adducts form between the ionizable lipid and RNA, a conclusion contradicting the reference study and the known literature. The AI Scientist-v2 from Sakana AI on the MD project claimed in its final paper to have run molecular dynamics simulations of hundreds of nanoseconds, whereas no simulations were run and a synthetic dataset with fabricated feature vectors was used instead. In all these cases, erroneous outputs were embedded in otherwise plausible-looking reports, making it nonevident whether checking AI results or redoing the project from scratch without AI would be more time-effective.

Certain categories of tasks proved systematically problematic, as we conclude from the analysis of the failure patterns. These tasks included identification of extrema (e.g., find the best-performing model or featurization), statements involving universal or existential quantification over a set of conditions, comparisons of results across methods or scenarios, and questions involving variables with ambiguous numerical definitions (e.g., germline closeness in the antibody project). These task types should be best treated with additional caution in the design of research prompts for AI frameworks.

We also identified cases of formally valid but scientifically misleading reasoning. For example, in the antibody developability project, one run reported that deep learning embeddings do not improve prediction over one-hot encoding, a conclusion matching the reference paper. The stated conclusion was technically correct but deeply misleading because it implied the same qualitative result as the original paper when in fact the performance was entirely different. Overconfidence was another recurring feature of the final reports. For instance, in the MD project, a detailed discussion of a tail ketone as a secondary interaction site was based on a single frame from a ten-frame trajectory covering only 10 ps of simulation. Such presentation could mislead users who lack specific domain expertise, further questioning the practical effectiveness of AI frameworks.

Overall, our present results confirm and extend the central conclusion of the first paper: real published research from leading pharmaceutical companies represents a substantially more difficult task for current AI frameworks than solving standard benchmarks or evaluation tasks specifically designed for AI testing. The two projects examined here, despite differing markedly in methodology, exhibited the same overall pattern of partial successes coexisting with severe failures, narrower scope, and substantial irreproducibility across runs. The fact that this pattern is reproduced across five distinct case studies from various scientific domains and five evaluated AI frameworks suggests that it reflects the current state of the field rather than the limitations of any particular framework or task. As in the first paper, we conclude that AI frameworks in their present form could be more useful for prototyping research directions and for stress-testing studies that have already been substantially completed by human researchers, rather than conducting autonomous scientific research.

## Methods

### Paper selection and Prompt engineering

Papers were selected to reflect the current frontier in the corresponding fields, with practical relevance as the primary criterion. For each project, a full-scale prompt was formulated describing the research problem and the key scientific questions, without prescribing specific datasets or methods. The full-scale prompt for the antibody project covered the scope of the entire reference paper, while for the lipid-RNA project it covered only the computational modeling component of the reference paper (see Appendix). Smaller-scope prompts were then designed to reduce the scope by roughly an order of magnitude: for the antibody project, the analysis was narrowed to two specific datasets, two prescribed sequence representations, and a fixed evaluation procedure; for the MD project, molecular structures, force fields, and protocols were explicitly prescribed, and only construction and short equilibration of three all-atom systems were requested, without production MD or its analysis (see Appendix).

For all other aspects of Methods, we refer the reader to the first paper in this series,^8^ We remind the reader that AI frameworks were run in January-March 2026, and therefore, our conclusions reflect the state of the listed AI frameworks at that time, which may have changed since then.^15,32^

## Acknowledgements

We are deeply grateful to Dr. Diego del Alamo, Dr. Yves F. Nanfack (Takeda Pharmaceuticals Research & Development), and Dr. Younghoon Oh (Eli Lilly and Company) for valuable discussions and detailed feedback on the manuscript. We would also like to thank the authors of Kosmos and K-Dense for providing free credits to students and academics to run these tools, and the authors of ToolUniverse, BioAgents, and the AI Scientist-v2 for publishing their code open-source and under permissive licenses.

## APPENDIX

Prompts and smaller-scope prompts that we used are given below.

### Prediction of Antibody Developability Properties

#### Initial Prompt

Problem: Evaluating AI Models for Predicting Therapeutic Antibody Developability Properties

Therapeutic antibody development requires candidates to possess favorable biophysical properties to succeed in manufacturing and clinical trials. Current antibody design methods are typically evaluated with native sequence or structure recovery, which does not provide a complete indication of therapeutic potential.

Run advanced scientific research to answer the following research questions:

1. Can pretrained protein AI models predict antibody developability properties in a zero-shot manner? Which models perform best for each property category?
2. Does fine-tuning protein AI model embeddings on labeled developability data improve prediction accuracy compared to zero-shot approaches? How much training data is required to achieve significant improvements?
3. Do protein AI models learn genuine biophysical fitness landscapes, or do their predictions primarily reflect sequence similarity to germline? Compare structure-informed ML models, sequence-only ML models, and physics-based models.
4. What recommendations can be given to consider when developing zero-shot and few-shot antibody developability models?

Use publicly available antibody developability datasets containing experimental measurements. Don’t use synthetic (computationally generated) data. Evaluate using Spearman’s rank correlation between model outputs and experimental values.

After conducting this research, write a research paper with introduction, results, discussion, methods, and references.

#### Smaller-Scope Prompt

Problem: Prediction of Antibody Developability Properties Using Protein Language Model Embeddings

Therapeutic antibody development requires candidates to possess favorable developability properties in order to succeed in manufacturing and clinical trials. Computational prediction of these properties can help prioritize antibody candidates earlier in the design process. Your task is to evaluate whether protein language model embeddings improve prediction of antibody developability properties compared with a simple sequence-based baseline representation.

Use two publicly available experimental antibody datasets that measure different developability properties: the antibody expression dataset reported by Adams et al. (2017), which contains antibody amino acid sequences and experimentally measured expression levels, and the antibody binding affinity dataset reported by Adams et al. (2017), which contains antibody amino acid sequences and experimentally measured binding affinity values. Both datasets are available in the FLAb benchmark repository https://github.com/Graylab/FLAb/tree/main/data The expression dataset can be accessed at: https://github.com/Graylab/FLAb/blob/main/data/expression/adams2017measuring_4420-fluorescein_exp_er.csv The binding dataset can be accessed at: https://github.com/Graylab/FLAb/blob/main/data/binding/adams2017measuring_4420-fluorescein_kd-titeseq.csv

Construct two types of sequence-derived feature representations. The first representation should consist of embeddings generated using the ESM-2 protein language model with 35M parameters also known as facebook/esm2_t12_35M_UR50D. The second representation should consist of a one-hot encoding of the antibody amino acid sequence. Represent each amino acid using a binary vector of length twenty corresponding to the twenty standard amino acids. For a sequence of length L, construct a matrix of size L x 20 in which each row contains a value of one at the position corresponding to the amino acid identity and zero in all other positions. Convert this matrix into a single feature vector before using it as input to the regression model.

For the expression dataset, predict the experimentally measured expression value reported in the dataset. For the binding dataset, predict negative log10 value of the experimentally measured binding affinity constant Kd.

Perform few-shot supervised learning by training downstream regression models using the sequence representations. Clearly describe the regression models used and explain how the embedding features or one-hot sequence features are used as model inputs.

Evaluate predictive performance using Spearman rank correlation between predicted values and experimental measurements. Use five-fold cross-validation with leakage-safe splits that prevent highly similar antibody sequences from appearing in both training and validation folds. Report the Spearman correlation for each fold and the mean performance across all folds. Don’t use synthetic (computationally generated) data instead of experimental ones. Calculate p-values wherever applicable.

The final report should clearly describe the datasets used, the developability properties evaluated, the sequence representations constructed, the regression models applied in the few-shot setting, the evaluation procedure, and the resulting predictive performance. The report should compare embedding-based models with the one-hot sequence baseline and discuss how representation choice and training data size influence predictive performance.

### Modeling Lipid-RNA Interactions with MD simulations

#### Initial Prompt

Problem: Investigating Non-covalent Interactions Between Oxidized Ionizable Lipids and siRNA in Lipid Nanoparticle Systems

Lipid nanoparticles (LNPs) are clinically approved delivery vehicles for siRNA therapeutics. The ionizable lipid DLin-MC3-DMA (MC3) is a key component of approved formulations but contains bis-allylic unsaturated hydrocarbon tails susceptible to oxidation. Oxidation of these tails can produce conjugated dienone byproducts containing ketone functional groups. Understanding how such oxidative modifications affect lipid-RNA interactions within LNPs is important for predicting drug product stability and performance.

Run all-atom molecular dynamics simulations to answer the following research questions:

1. Does the introduction of a ketone group into the MC3 lipid tail through oxidation create new hydrogen bonding interactions with encapsulated siRNA? If so, how do these tail-mediated interactions compare in magnitude to the established interactions between the ionizable amine headgroup and RNA?
2. How do the radial distribution functions characterizing lipid-RNA proximity differ between native MC3 and its oxidized dienone byproduct?
3. For siRNA containing 2’-OH groups (unmodified ribose), does the oxidized lipid species show additional interaction modes compared to fully 2’-O-methyl modified siRNA?

The system should include double-stranded siRNA with chemically modified backbones (phosphorothioate linkages) and sugar modifications (2’-O-methyl and 2’-fluoro), protonated ionizable lipid species, and aqueous solvent conditions representative of physiological buffer. Two separate simulation systems should be constructed for comparison: one with native MC3 molecules, and the other with oxidized MC3 dienone byproduct. The siRNA sequence to be used in this study is: sense strand 5’-uscscuauGfaCfUfGfua-gauuuusasu-3’ and antisense strand 5’-pasUfsaaaAfucuacagUfcA-fuaggasasu-3’. The chemical modifications are denoted as follows: a, c, g, u: 2’-O-Methyl nucleotides, s: phosphorothioate; Af, Cf, Gf, Uf: 2’-Fluoronucleotides, p: (mono)phosphate.

After conducting this research, write a research paper with introduction, results, discussion, methods, and references.

#### Smaller-Scope Prompt

Problem: Modeling Non-covalent Interactions Between Oxidized Ionizable Lipids and siRNA in Lipid Nanoparticle Systems

Lipid nanoparticles (LNPs) are clinically approved delivery vehicles for siRNA therapeutics. The ionizable lipid DLin-MC3-DMA (MC3) is a key component of approved formulations but contains bis-allylic unsaturated hydrocarbon tails susceptible to oxidation. Oxidation of these tails can produce conjugated dienone byproducts containing ketone functional groups. Understanding how such oxidative modifications affect lipid-RNA interactions within LNPs is important for predicting drug product stability and performance.

Retrieve native DLin-MC3-DMA from PubChem (CID 49785164). Construct the oxidized MC3 dienone byproduct with E,Z-conjugated dienone geometry in one tail. Report the SMILES strings and 2D structural diagrams for both from the computed files directly, not from memory.

Builld an all-atom model of a molecular system that includes double-stranded siRNA with chemically modified backbones (phosphorothioate linkages) and sugar modifications (2’-O-methyl and 2’-fluoro), protonated native MC3 species, and aqueous solvent conditions representative of physiological buffer in a cubic simulation box with each side set to 9 nm. The siRNA sequence to be used in this study is: sense strand 5’-uscscuauGfaCfUfGfua-gauuuusasu-3’ and antisense strand 5’-pasUfsaaaAfucuacagUfcA-fuaggasasu-3’. The chemical modifications are denoted as follows: a, c, g, u: 2’-O-Methyl nucleotides, s: phosphorothioate; Af, Cf, Gf, Uf: 2’-Fluoronucleotides, p: (mono)phosphate.

In a similar way, build a second system with the same siRNA and oxidized MC3, and a third system with oxidized MC3 and with siRNA containing 2’-OH groups (unmodified ribose).

For each of these three systems, perform energy minimization and 1 ns NPT equilibration. Use the CHARMM36 force field, including nucleic acids and its extension for modified nucleotides, to model the modified siRNA, and the CHARMM General Force Field to parameterize the cationic ionizable lipids. Make sure the systems stay stable, otherwise rebuild them.

After conducting this research, write a research paper with introduction, results, discussion, methods, and references.

## References

1. Phan L, Gatti A, Li N, et al. Humanity’s Last Exam. arXiv. 2025; doi: 10.48550/arXiv.2501.14249

2. Phan L, Gatti A, Li N, et al. A benchmark of expert-level academic questions to assess AI capabilities. Nature. 2026; 649(8099):1139–1146. doi:10.1038/s41586-025-09962-4

3. Rein D, Hou BL, Stickland AC, et al. GPQA: A Graduate-Level Google-Proof Q&A Benchmark. arXiv. 2023; doi: 10.48550/arXiv.2311.12022

4. Artificial Analysis. https://artificialanalysis.ai/

5. Kapoor S, Kirgis P, Schwartz A, et al. Open-World Evaluations for Measuring Frontier AI Capabilities. arXiv. 2026; doi: 10.48550/arXiv.2605.20520

6. Agrawal S, Anadkat HB, Athimoolam KK, et al. Can AI Conduct Autonomous Scientific Research? Case Studies on Two Real-World Tasks. bioRxiv. 2026; doi: 10.64898/2026.01.05.697809

7. Gulluoglu HSA, Baby J, Bagul KM, et al. Practical Use of Advanced AI Frameworks on Real-Life Scientific Problems: Three Case Studies. bioRxiv. 2026; doi: 10.64898/2026.06.23.734132

8. Ahmed MO, Amale SA, Bhavsar RD, et al. Real Science Is Harder Than Benchmarks: Evaluating Advanced AI Frameworks on Published Studies. I. Uncertainty Quantification, ML on Therapeutic Data Commons, and Agent-Based Modeling. bioRxiv. 2026; doi: 10.64898/2026.06.24.734302

9. Kirgis P, Kapoor S, Schwartz A, et al. Can AI agents conduct open-ended AI research? Early evidence from two case studies. arXiv. 2026; doi: 10.48550/arXiv.2607.27191

10. Jiang H, Chase J, Fu L, Shivaram S, Sinitskiy AV. Case Study of Using AI as Co-Pilot in Biotech Research: Functional Network Analysis of Invasive Cancer. bioRxiv. 2025; doi: 10.1101/2025.05.14.654152

11. Kirgis P, Kapoor S, Rabanser S, et al. Log analysis is necessary for credible evaluation of AI agents. arXiv. 2026; doi: 10.48550/arXiv.2605.08545

12. Chungyoun M, Gray J. Fitness Landscape for Antibodies 2: Benchmarking Reveals That Protein AI Models Cannot Yet Consistently Predict Developability Properties. bioRxiv. 2025; doi:10.64898/2025.12.27.696706

13. Raybould MIJ, Marks C, Krawczyk K, et al. The Therapeutic Antibody Profiler (TAP): Five Computational Developability Guidelines. bioRxiv. 2018; doi:10.1101/359141

14. Tan P, Li S, Huang J, Zhou Z, Hong L. Harnessing deep learning to accelerate the development of antibodies and aptamers. Acta Pharm Sin B. 2026; 16(2):788–801. doi:10.1016/j.apsb.2025.12.017

15. Arora R, Chen LT, D. M, Marks DS, Church GM. PG-LLM: Benchmarking General-Purpose Language Models for Protein Variant Ranking. bioRxiv. 2026; doi: 10.64898/2026.07.27.741045

16. Wang T, He X, Li M, et al. Ab initio characterization of protein molecular dynamics with AI(2)BMD. Nature. 2024; 635(8040): 1019–1027. doi:10.1038/s41586-024-08127-z

17. Costa AdS, Mitnikov I, Pellegrini F, et al. EquiJump: Protein Dynamics Simulation via SO(3)-Equivariant Stochastic Interpolants. arXiv. 2024; doi: 10.48550/arXiv.2410.09667

18. Batatia I, Kovács DP, Simm GNC, Ortner C, Csányi G. MACE: Higher Order Equivariant Message Passing Neural Networks for Fast and Accurate Force Fields. arXiv. 2022; doi: 10.48550/arXiv.2206.07697

19. Chigaev M, Smith JS, Anaya S, et al. Lightweight and effective tensor sensitivity for atomistic neural networks. J Chem Phys. 2023; 158(18); doi:10.1063/5.0142127

20. Zhang L, Han J, Wang H, Car R E W. Deep Potential Molecular Dynamics: A Scalable Model with the Accuracy of Quantum Mechanics. Phys Rev Lett. 2018; 120(14):143001. doi:10.1103/PhysRevLett.120.143001

21. Wang H, Zhang L, Han J E W. DeePMD-kit: A deep learning package for many-body potential energy representation and molecular dynamics. Computer Physics Communications. 2018; 228:178–184. doi:10.1016/j.cpc.2018.03.016

22. Smith JS, Bettencourt M, Pellegrini F, Glines F, Kucukbenli E. Enabling Scalable AI-Driven Molecular Dynamics Simulations. https://developer.nvidia.com/blog/enabling-scalable-ai-driven-molecular-dynamics-simulations/

23. Jia W, Wang H, Chen M, et al. Pushing the limit of molecular dynamics with ab initio accuracy to 100 million atoms with machine learning. arXiv. 2020; doi: 10.48550/arXiv.2005.00223

24. Lu D, Wang H, Chen M, et al. 86 PFLOPS Deep Potential Molecular Dynamics simulation of 100 million atoms with ab initio accuracy. Computer Physics Communications. 2021; 259:107624. doi:10.1016/j.cpc.2020.107624

25. Moore H. Biomolecular Simulations with Machine Learning Potentials (PhD thesis). University of Cambridge, 2025.

26. Kovacs DP, Moore JH, Browning NJ, et al. MACE-OFF: Short-Range Transferable Machine Learning Force Fields for Organic Molecules. J Am Chem Soc. 2025; 147(21):17598–17611. doi:10.1021/jacs.4c07099

27. Chandrasekhar A, Farimani AB. Automating MD simulations for Proteins using Large language Models: NAMD-Agent. arXiv. 2025; doi: 10.48550/arXiv.2507.0788

28. Campbell Q, Cox S, Medina J, Watterson B, White AD. MDCrow: Automating Molecular Dynamics Workflows with Large Language Models. arXiv. 2025; doi: 10.48550/arXiv.2502.09565

29. Jain T, Sun T, Durand S, et al. Biophysical properties of the clinical-stage antibody landscape. Proc Natl Acad Sci U S A. 2017; 114(5):944–949; doi:10.1073/pnas.1616408114

30. https://huggingface.co/datasets/ginkgo-datapoints/GDPa1

31. Estabrook DA, Huang L, Lucchese OR, et al. Buffer optimization of siRNA-lipid nanoparticles mitigates lipid oxidation and RNA-lipid adduct formation. Nat Commun. 2025; 16(1):8380. doi:10.1038/s41467-025-63651-4

32. Shanehsazzadeh A. Autonomous de novo protein binder design with Claude. 2026. https://www.alphaxiv.org/abs/2608.claude-de-novo

